# Novel VCP activator reverses multisystem proteinopathy nuclear proteostasis defects and enhances TDP-43 aggregate clearance

**DOI:** 10.1101/2023.03.15.532082

**Authors:** Jessica M. Phan, Benjamin C. Creekmore, Aivi T. Nguyen, Darya D. Bershadskaya, Nabil F. Darwich, Edward B. Lee

**Author notes:** **Correspondence to:** Edward B. Lee, 613A Stellar Chance Laboratories, 422 Curie Blvd, Philadelphia, PA 19104. Equal contributions (Jessica M. Phan, Benjamin C. Creekmore).

## Abstract

Pathogenic variants in *VCP* cause multisystem proteinopathy (MSP), a disease characterized by multiple clinical phenotypes including inclusion body myopathy, Paget’s disease of the bone, and frontotemporal dementia (FTD). How such diverse phenotypes are driven by pathogenic VCP variants is not known. We found that these diseases exhibit a common pathologic feature, ubiquitinated intranuclear inclusions affecting myocytes, osteoclasts and neurons. Moreover, knock-in cell lines harboring MSP variants show a reduction in nuclear VCP. Given that MSP is associated with neuronal intranuclear inclusions comprised of TDP-43 protein, we developed a cellular model whereby proteostatic stress results in the formation of insoluble intranuclear TDP-43 aggregates. Consistent with a loss of nuclear VCP function, cells harboring MSP variants or cells treated with VCP inhibitor exhibited decreased clearance of insoluble intranuclear TDP-43 aggregates. Moreover, we identified four novel compounds that activate VCP primarily by increasing D2 ATPase activity whereby pharmacologic VCP activation appears to enhance clearance of insoluble intranuclear TDP-43 aggregate. Our findings suggest that VCP function is important for nuclear protein homeostasis, that MSP may be the result of impaired nuclear proteostasis, and that VCP activation may be potential therapeutic by virtue of enhancing the clearance of intranuclear protein aggregates.

## Introduction

Inclusion body myopathy associated with Paget disease of bone and frontotemporal dementia (IBMPFD, also known as multisystem proteinopathy, MSP) is a fatal disease that affects multiple organ systems including muscle, bone, and/or brain and causes progressive muscle weakness, defects in bone formation, and/or frontotemporal dementia (FTD) (1,2). MSP is caused by pathogenic variants affecting valosin-containing protein (VCP), a member of the ATPases associated with diverse cellular activities (AAA+) superfamily of proteins (3,4).

The structural conformation of VCP and the coordination of its subunits are critical for its role in substrate processing (5). VCP is made up of an N-terminal domain and two ATPase domains, D1 and D2. Characteristic of other AAA+ proteins, VCP forms a homohexameric structure with a central pore. VCP, together with cofactors such as UFD1 and NPLOC4 (UN), bind to poly-ubiquitinated substrates and unfolds ubiquitin followed by the attached substrate by pulling the peptide chain through its central pore (6–8). Recently, we reported that VCP also has activity against poly-ubiquitinated protein aggregates in vitro (9). During this process of protein unfolding, the D1 and D2 ATPase domains of VCP seem to have distinct roles in substrate processing. The D1 ATPase domain needs to bind, but not hydrolyze ATP for VCP to unfold substrate. The D2 ATPase domain, however, uses ATP hydrolysis to processively pull unfolded substrate through VCP’s central pore (8,10).

VCP is a highly conserved protein and is involved in various cellular pathways including autophagy, membrane fusion, and cell cycle control where VCP is typically involved in extracting or unfolding key protein substrates to affect downstream organelle function (11). Indeed, VCP is also critical in several protein control quality pathways (the ubiquitin proteasome system, endoplasmic-reticulum associated degradation, mitochondria-associated degradation, ribosomal associated degradation) in which VCP binds to poly-ubiquitinated substrates and utilizes mechanical energy driven by ATP hydrolysis to unfold ubiquitin and the attached substrates (6–8). Once unfolded by VCP, these substrates are then often targeted to the proteasome for degradation (12–14).

Although MSP can affect multiple and different organ systems, it is unclear whether there is a common pathology across muscle, bone, and brain tissue in the diseases associated with pathogenic VCP variants (2). The neuropathology of FTD in the setting of MSP is frontotemporal lobar degeneration with TDP-43 inclusions (FTLD-TDP) type D where ubiquitinated TDP-43 inclusions are found predominantly inside nuclei of affected neurons (15,16). Intranuclear inclusions are also a feature of inclusion body myositis (17,18). Structurally, pathogenic VCP variants associated with MSP are located near the interface of the N-terminal and D1 ATPase domains (19). In vitro biochemistry has shown these variants are associated with increased ATPase activity and enhanced substrate processing/unfolding (19). They also affect N-terminal domain structural dynamics and cofactor binding (19). Altered VCP function can lead to a variety of cellular defects as MSP variants have been associated with altered endoplasmic reticulum-associated degradation, autophagosome/lysosome maturation, endolysosomal processing, and mitochondrial fusion (11). We and others have shown that VCP appears to exhibit disaggregase activity against ubiquitinated inclusions thereby reducing intracellular protein aggregation (9,20–22). Thus, an important question is how pathogenic MSP variants in VCP, which are overactive in vitro, can lead to protein aggregation in vivo.

Transgenic and knock-in mouse models expressing pathogenic MSP variants have been reported to recapitulate multiple MSP disease phenotypes including skeletal muscle abnormalities, behavioral defects, and increased expression of ubiquitinated proteins while also being associated with cytoplasmic TDP-43 accumulation (22). While pathogenic VCP variants drive MSP disease pathogenesis, relatively few studies, mainly limited to yeast, have investigated VCP’s role in nuclear protein homeostasis (23,24). With evidence of pathological overlap of intranuclear inclusions across muscle and brain tissues, we wanted to explore VCP’s role in nuclear protein homeostasis.

In this study, we performed immunohistochemistry on FTLD-TDP, inclusion body myopathy, and Paget’s disease of bone tissue samples and identified the presence of ubiquitinated intranuclear inclusions in all three diseases linked to MSP. We then generated homozygous knock-in of pathogenic VCP variants into HeLa cells and found that cells harboring A232E and R155H VCP exhibited a reduction in nuclear VCP expression. To test VCP’s effects on TDP-43 protein homeostasis in mutant cells, we modeled nuclear TDP-43 aggregation by utilizing a construct of TDP-43 deficient in RNA binding (TDP-4FL) that, when expressed in cells, forms soluble nuclear inclusions. We found that when the proteasome is blocked in cells expressing TDP-4FL, these intranuclear inclusions became larger, ubiquitinated and insoluble. When TDP-4FL was expressed in cells expressing VCP-A232E and R155H variants, or VCP was inhibited with CB5083, a specific VCP inhibitor, clearance of TDP-4FL aggregates was disrupted. Additionally, we identified 4 novel small molecule VCP activators that stimulate VCP’s D2 ATPase activity. Moreover, when cells were treated with VCP activator, clearance of TDP-4FL aggregates was augmented. These results suggest that nuclear reduction of VCP expression associated with pathogenic MSP variants results in a loss of nuclear protein homeostasis and reduced clearance of intranuclear TDP-43 aggregates, and that this phenotype may be rescued by a small molecule VCP activator.

## Result

### Ubiquitinated intranuclear inclusions in FTLD-TDP type D, inclusion body myopathy and Paget’s disease of bone

Given the diverse tissues that are affected in multisystem proteinopathy, we sought to identify whether there is a common pathology across FTLD-TDP type D, inclusion body myopathy, and Paget’s disease of bone. In individuals with frontotemporal dementia, pathogenic VCP variants are known to be associated with ubiquitinated neuronal intranuclear inclusions comprised of TDP-43 protein (representative examples shown in Figure 1A) (25,26). Similarly, inclusion body myopathy is known to be associated with ubiquitin-positive intranuclear inclusions consisting of an unknown protein, as shown by electron microscopy and immunohistochemistry (Figure 1B) (17,27,28). Whether Paget’s disease of bone, a disease associated with overactive osteoclasts, also exhibits ubiquitinated intranuclear inclusions has not been determined. Ten bone biopsies from cases of Paget’s disease of bone were immunostained for ubiquitin, of which two biopsies were excluded due to scant tissue with rare to absent osteoclasts. The remaining eight cases showed osteoclasts with strongly ubiquitin-positive inclusions including conspicuous, spherical intranuclear inclusions together with a few granular, apparently cytoplasmic aggregates (Figure 1C). Osteoclasts from normal bone (n=10) or other osteoclast rich lesions including giant cell tumor of bone (n=10) and fracture calluses (n=10) did not exhibit intranuclear inclusions. Our findings are consistent with prior electron microscopy studies indicating that osteoclasts in Paget’s disease of bone exhibit abnormal intranuclear inclusions (18), and suggests that these three disparate diseases associated with pathogenic VCP variants share a common pathology.

**Figure 1:**
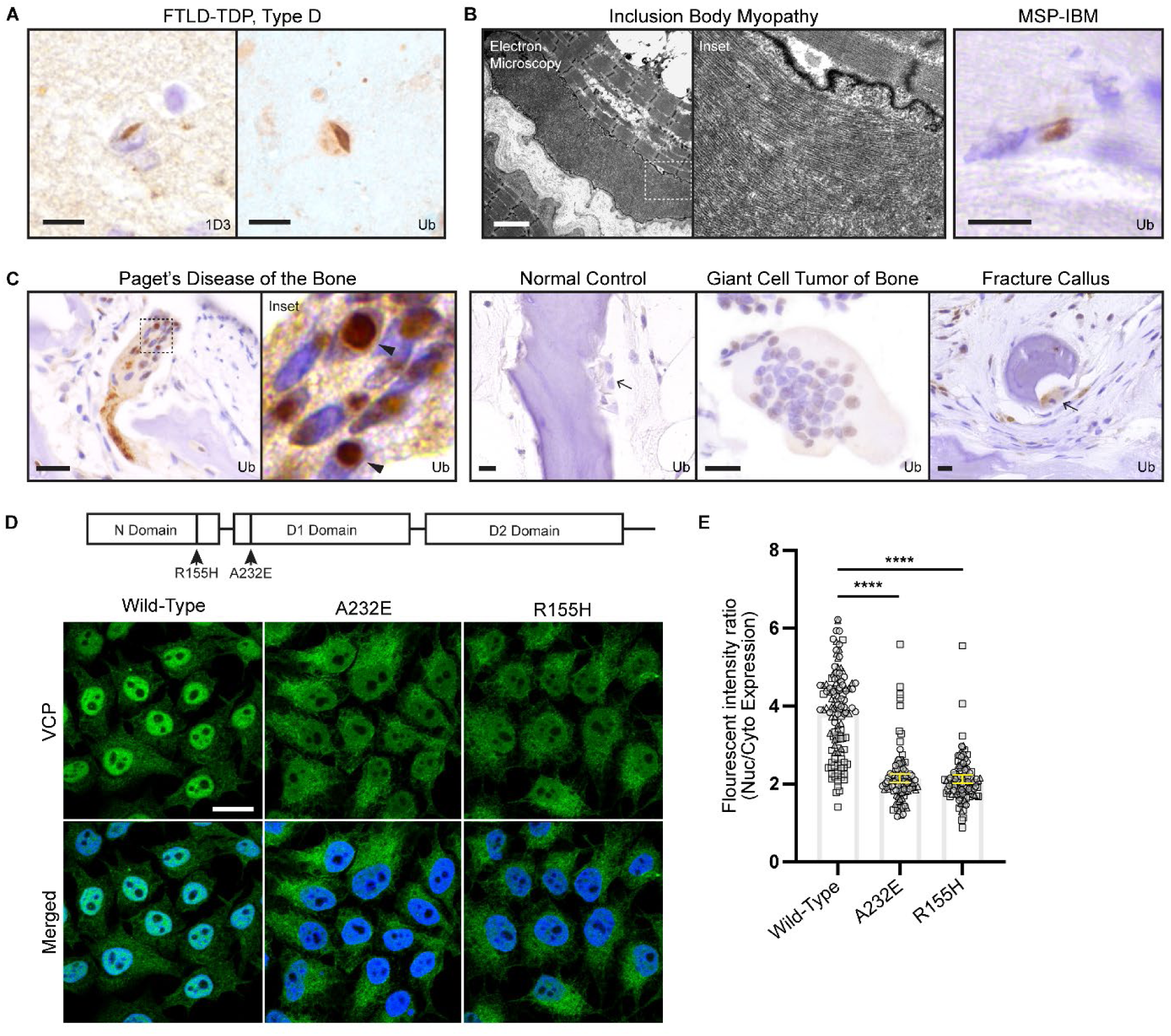
Ubiquitinated intranuclear inclusions in FTLD-TDP, IBM and Paget’s disease of bone, and reduced nuclear VCP localization in CRISPR edited cells harboring pathogenic VCP variants. **(A)** Immunohistochemical stains of FTLD-TDP type D neocortical brain tissue for phosphorylated TDP-43 (1D3) or ubiquitin. Scale bar, 10µm. **(B)** Electron microscopy and immunohistochemical stain for ubiquitin of inclusion body myopathy muscle. Scale bars: electron microscopy 2µm; immunohistochemistry 10µm. **(C)** Immunohistochemical stains for ubiquitin of Paget’s disease bone, normal control bone, giant cell tumor of the bone, or fracture callus. Arrowheads point to ubiquitin-positive intranuclear inclusions and arrows point to osteoclasts without intranuclear inclusions. Scale bars, 10µm. **(D)** Confocal images of immunofluorescence staining for VCP (green) and DAPI (blue) of WT HeLa cell lines versus CRISPR-Cas9 knock-in HeLa cell lines harboring A232E or R155H VCP pathogenic variants. Scale bar, 15µm. **(E)** Ratio of nuclear to cytoplasmic VCP immunofluorescence intensity (n = 325 cells across all three cell lines analyzed from 3 independent experiments; data shown as individual data points with overall beta ± SE. Linear mixed effect model, **** P < 0.0001).

### Nuclear VCP expression is reduced in HeLa cells expressing A232E and R155H variants

The presence of ubiquitin-positive intranuclear inclusions in FTLD-TDP type D, inclusion body myopathy, and Paget’s disease of bone suggested that these diseases are associated with a loss of nuclear proteostasis. Prior overexpression studies have suggested that pathogenic VCP variants are partially excluded from the nucleus (29,30). To determine whether pathogenic VCP variants result in changes in the subcellular localization of endogenous VCP, CRISPR-Cas9 was used to edit the endogenous *VCP* locus by knocking-in two different pathogenic *VCP* variants into HeLa cells: p.A232E (the variant associated with the highest ATPase activity in vitro) and p.R155H (the most common pathogenic MSP variant). Targeted sequencing of knock-in cell lines at the top predicted off-target genomic sites failed to reveal off-target mutations (see Supplemental Table 1). Mutant cell lines were immunostained for VCP and compared to parental HeLa cells which revealed that VCP is localized to both the cytoplasm and nucleus. However, A232E or R155H knock-in cell lines showed a reduction in the relative amount of nuclear VCP (Figure 1D). Quantitative image analysis showed that there is approximately a 60% reduction in the fluorescence intensity ratio of nuclear to cytoplasmic VCP signal in A232E and R155H cells compared to wild-type cells (P < 0.0001; Figure 1E).

### Proteasome inhibition promotes the formation of insoluble ubiquitinated intranuclear TDP-43 inclusions

We hypothesized that the reduced nuclear localization of mutant VCP would lead to diminished nuclear proteostasis. To better study VCP’s role in nuclear protein homeostasis, we focused our studies on TDP-43 since it is known to be the protein that forms inclusions in FTLD-TDP type D brain tissue. First, we expressed myc-tagged TDP-43 protein which showed diffuse nuclear expression in wild-type cells (Supplemental Figure 1A). To promote the formation of intranuclear TDP-43 inclusions, cells were treated with MG132 to inhibit the proteasome which resulted in the formation of coarse nuclear punctae as demonstrated by immunofluorescence confocal microscopy (Supplemental Figure 1A). Moreover, these nuclear aggregates were positive for both ubiquitin and VCP (Supplemental Figure 1E, 1F). These morphologic changes were associated with a reduction in soluble TDP-43 protein and an increase in insoluble TDP-43 protein over time as determine by immunoblotting (P = 0.170 for soluble protein; P < 0.05 for insoluble protein; Supplemental Figure 1B, 1C, 1D).

RNA binding has been demonstrated to alter TDP-43 dynamics, promoting self-association into phase separated puncta called anisosomes (31–34). We therefore tested whether an RNA-binding deficient TDP-43 mutant containing phenylalanine to leucine mutations within RNA recognition motifs 1 and 2 (F147/149/229/231L, hereafter referred to as TDP-4FL) would result in an even more robust model of intranuclear TDP-43 inclusions (32). Expression of TDP-4FL led to the formation of small, intranuclear, phase-separated anisosomes, as reported previously (32,33). Moreover, when the proteasome was inhibited with MG132 in cells expressing TDP-4FL, the size, morphology, and solubility of TDP-4FL protein inclusions were altered. At 3 or 6 hours of MG132 treatment, the small anisosome structures coalesced into larger, irregularly shaped aggregates. Confocal double immunofluorescence images of MG132-treated cells revealed that these TDP-4FL aggregates co-localize with ubiquitin (Figure 2A) and VCP (Figure 2B). To test the solubility of the TDP-4FL aggregates, sequential biochemical extraction of cells were used to separate detergent (RIPA) soluble proteins versus RIPA insoluble (urea soluble) proteins. MG132 treatment over time resulted in a decrease in soluble TDP-4FL (Figure 2C, 2D) with a concomitant increase in insoluble TDP-4FL (P < 0.01 for soluble protein; P < 0.05 for insoluble protein; Figure 2C, 2E). Immunoprecipitation of insoluble myc-TDP-4FL also revealed that MG132 treatment resulted in ubiquitination of TDP-4FL protein (Figure 2F).

**Figure 2:**
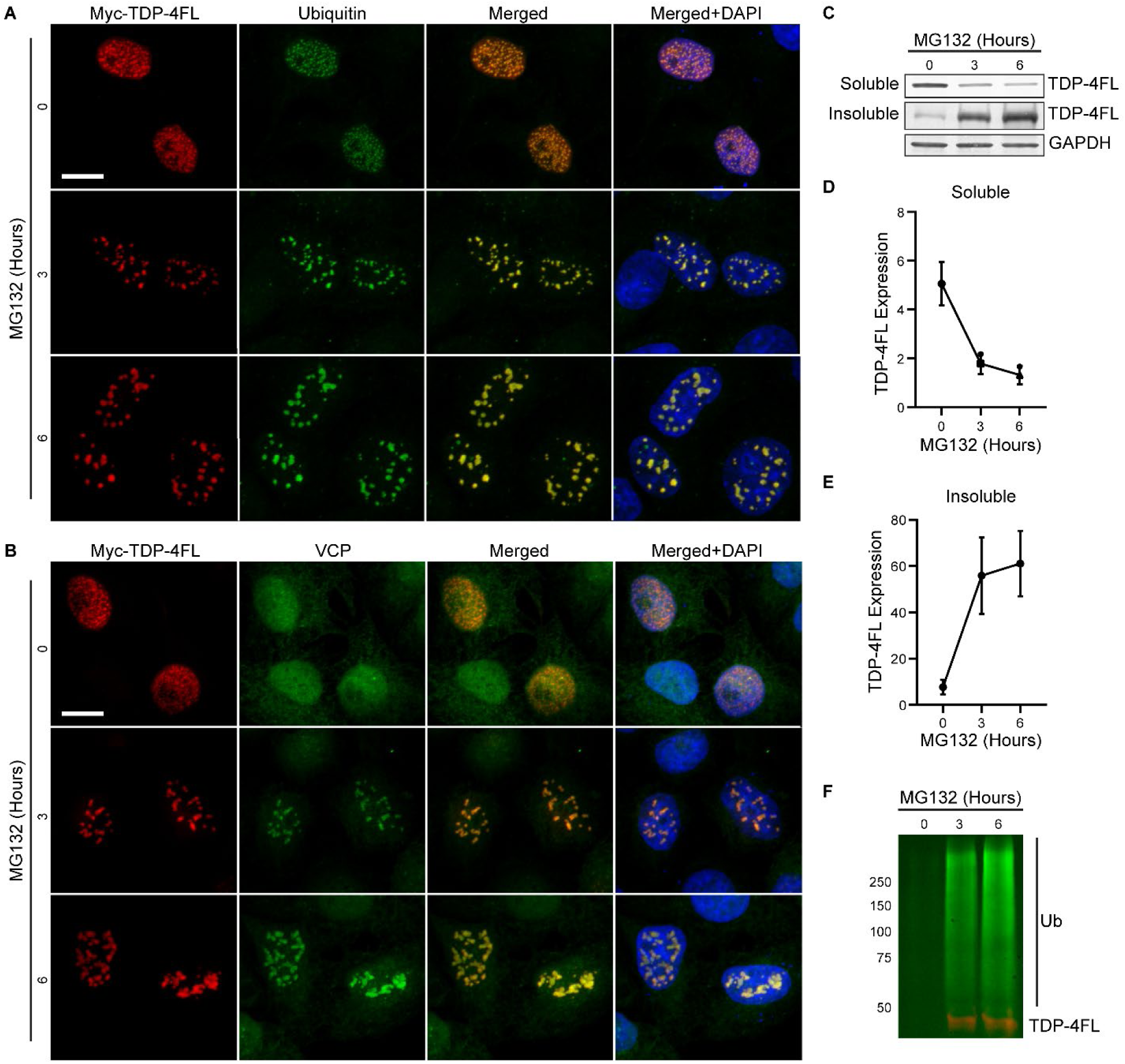
Proteasome inhibition promotes insoluble intranuclear TDP-4FL inclusions that co-localize with ubiquitin and VCP. **(A, B)** Immunofluorescence confocal images of HeLa cells expressing TDP-4FL treated with 4μM MG132 for 0, 3, or 6 hours stained for myc (TDP-4FL, red) and either ubiquitin (A, green) or VCP (B, green). Scale bar, 10µm. **(C)** Immunoblots for myc (TDP-4FL) from soluble and insoluble protein fractions from HeLa cells expressing TDP-4FL treated with 4μM MG132 for 0, 3, or 6 hours. GAPDH provided as a loading control. **(D-E)** Quantification of (D) soluble or (E) insoluble TDP-4FL immunoblots. (n = 4 independent experiment, results are expressed as mean ± SEM over time. One-way ANOVA, ** P < 0.01 for soluble protein, *P < 0.05 for insoluble protein). **(F)** Anti-myc immunoprecipitation from insoluble protein fractions, immunoblotted for myc (TDP-4FL, red) and ubiquitin (green).

### VCP variants lead to a defect in nuclear protein homeostasis

Because TDP-4FL inclusions co-localized with ubiquitin and VCP, we hypothesized that VCP may help maintain nuclear protein homeostasis and play a role in the clearance of TDP-4FL aggregates. To assess VCP’s effects on TDP-4FL protein homeostasis in mutant cells, we over-expressed TDP-4FL in our VCP-A232E and R155H cell lines. Interestingly, a higher proportion of A232E and R155H cells expressed TDP-4FL compared to wild-type cells (P < 0.01; Dunnett’s multiple comparison post-hoc P < 0.05, P < 0.01; Figure 3A). This result was not due to altered transfection efficiency as there was no difference in the proportion of cells expressing a control GFP construct across cell lines, suggesting that increased TDP-4FL expression in the mutant cells may be driven by defects in VCP function (Figure 3B).

**Figure 3:**
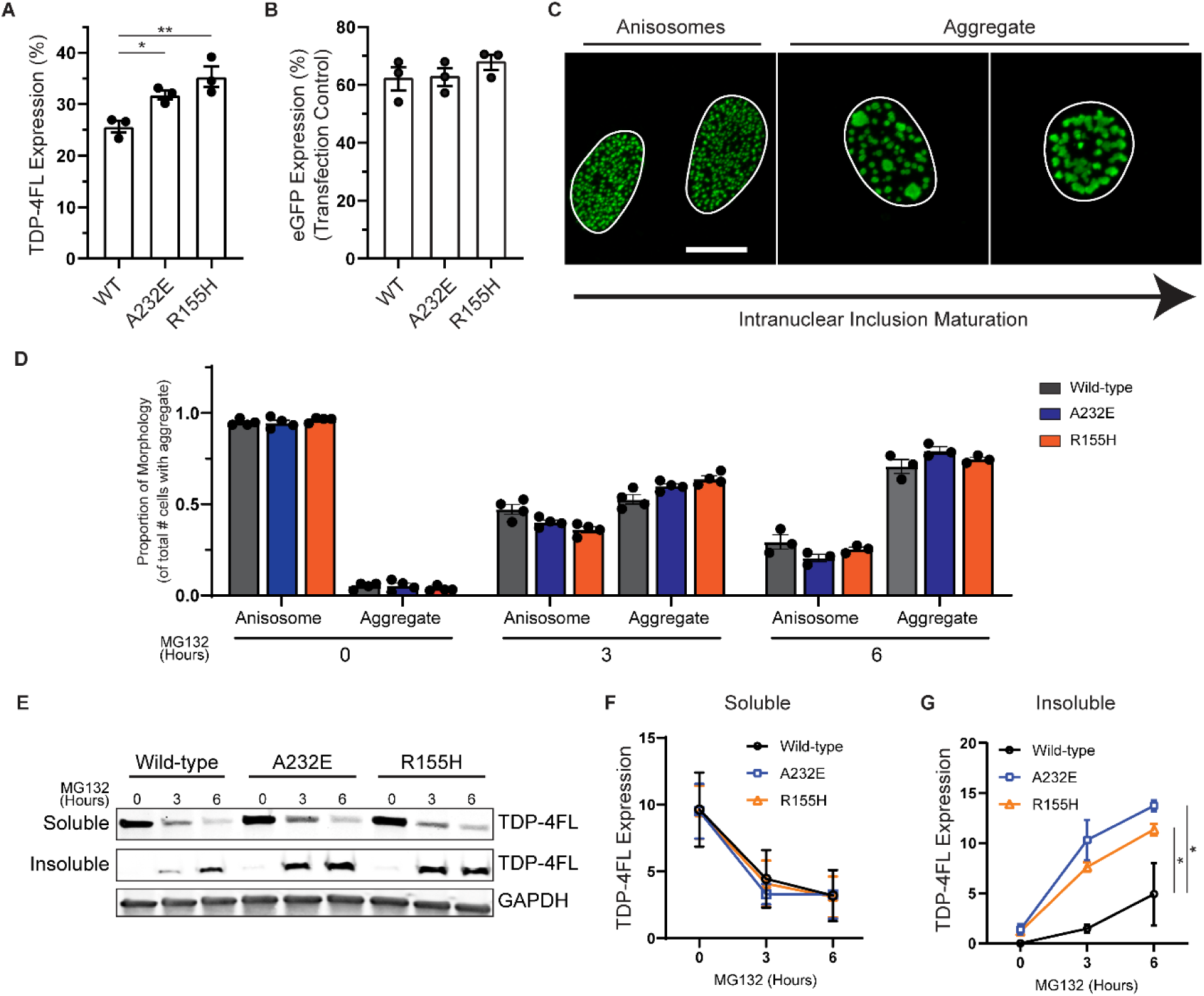
TDP-4FL anisosomes become larger and more insoluble due to pathogenic A232E and R155H VCP variants. **(A-B)** Analysis of immunofluorescence images to determine the percentage of cells expressing **(A)** TDP-4FL or **(B)** eGFP as a percentage of total cell counts after 24 hours of transient TDP-4FL or eGFP transfection in wild-type (WT) HeLa cells or CRISPR-Cas9 knock-in HeLa cells harboring A232E and R155H VCP pathogenic variants. (TDP-4FL, n = 6390 cells counted across all three cell lines over 3 independent experiments; results are expressed as mean percentage values from each independent experiment in addition to overall mean ± SEM. One-way ANOVA, **P < 0.01; Dunnett’s multiple comparison post-hoc, *P < 0.05, **P < 0.01. eGFP, n = 2135 cells counted across all three cell lines over 3 independent experiments; results are expressed as mean percentage values from each independent experiment in addition to overall mean ± SEM. One-way ANOVA, P = 0.459) **(C)** Representative confocal images of immunofluorescence for TDP-4FL with no MG132 treatment or 4 µM MG132 for 6 hours. Nuclear structures were categorized as either anisosomes or aggregates based on morphology. Scale bar, 10µm. **(D)** Analysis of immunofluorescence images to determine the proportion of anisosome versus aggregate morphology in WT, A232E, or R155H cells treated with 0, 3, or 6 hours of 4 µM MG132 (n = 7058 cells counted across all three cell lines over 3 independent replicates; results are expressed as mean proportion values from each independent experiment in addition to overall beta ± SE, linear mixed effects model for A232E cells: time P < 0.0001, genotype P = 0.708, time x genotype **P < 0.01; linear mixed effects model for R155H cells: time P < 0.0001, genotype P = 0. 734, time x genotype *P < 0.05). **(E)** Immunoblots of soluble and insoluble TDP-FL protein in WT, A232E, and R155H cells treated with 0, 3, 6 hours of 4µM MG132. GAPDH shown as a loading control. **(F-G)** Quantification of immunoblots for (F) soluble and (G) insoluble TDP-FL protein in WT, A232E, and R155H cells treated with 0, 3, 6 hours of 4 µM MG132. (n = 3, results expressed as mean ± SEM over time; two-way ANOVA for soluble protein: time **P < 0.01, genotype P = 0.690, time x genotype P = 0.513; for insoluble protein: time **P < 0.01, genotype *P < 0.05, time x genotype P = 0.210; Dunnett’s post-hoc analysis: *P <0.05).

Given that proteasome inhibition resulted in a change in the morphology of TDP-4FL inclusions from small anisosome to larger insoluble aggregate (Figure 3C), we assessed whether TDP-4FL inclusion morphology was altered in wild-type, A232E, and R155H cell lines with MG132 treatment over time. To test this, we analyzed 7058 TDP-4FL positive cells across all three cell lines over 3 independent replicates and categorized them in a blinded manner as having either small anisosomes or larger aggregates. In all cell lines, there was a reduction in anisosome morphology with an increase in aggregate morphology upon treatment with MG132 over time up to six hours. Moreover, there was also a small but statistically significant difference between cell lines, with both A232E and R155H cells exhibiting a higher proportion of cells with larger aggregate morphology (A232E genotype x time P < 0.01; R155H genotype x time P < 0.05; Figure 3D).

Given the relatively subtle shift in inclusion morphology associated with pathogenic VCP variants, TDP-4FL solubility was assessed across cell lines to determine whether biochemical analysis provided a more robust readout of intranuclear inclusion insolubility. Cells were extracted with RIPA buffer followed by a more stringent urea buffer in order to obtain soluble versus insoluble protein fractions. WT, A232E and R155H cells exhibited a similar decrease in soluble TDP-4FL protein levels with MG132 treatment over time (time P < 0.01, genotype P = 0.690, time x genotype P = 0.513; Figure 3E, 3F). However, at both 3 and 6 hours of MG132 treatment, insoluble TDP-4FL protein levels were significantly higher in A232E and R155H cells compared to WT cells (time P < 0.01, genotype P < 0.05, time x genotype P = 0.210; Dunnett’s post-hoc analysis P <0.05; Figure 3E, 3G).

### Clearance of insoluble intranuclear TDP-4FL aggregates is diminished by the VCP inhibitor CB5083

To examine the clearance and turnover of intranuclear TDP-4FL aggregates in a more dynamic manner, HeLa cells expressing TDP-4FL were treated with MG132 for three hours to induce formation of insoluble intranuclear aggregates followed by removal of MG132 and addition of the translational inhibitor cycloheximide (CHX) for 6 hours. This CHX recovery period allowed cells to begin to clear intranuclear TDP-4FL aggregates in the absence of any potential effect of new protein synthesis, using sequential biochemical fractionation followed by immunoblotting of TDP-4FL from soluble and insoluble protein fractions as a measure of intranuclear aggregate clearance (Figure 4A). This CHX recovery was done with or without the specific VCP inhibitor, CB5083, to determine whether VCP activity was involved in intranuclear aggregate clearance (Figure 4A). As described above, cells treated with 3 hours of MG132 exhibited an accumulation of larger, irregularly shaped, intranuclear TDP-4FL aggregates. Upon recovery in the presence of CHX, intranuclear aggregates were resolved with cells exhibiting small TDP-4FL anisosomes. However, co-treatment with both CHX and CB5083 resulted in residual irregularly shaped intranuclear TDP-4FL aggregates (Figure 4B). To provide biochemical evidence that VCP inhibition reduces intranuclear aggregate clearance, soluble and insoluble protein fractions were immunoblotted for TDP-4FL. While the CHX recovery resulted in a small increase in soluble TDP-4FL levels, the presence of CB5083 during the CHX recovery period was associated with a significant decrease in soluble TDP-4FL protein (P < 0.0001; Tukey’s post-hoc analysis P <0.01, P <0.001; Figure 4C, 4D). Even more evident was the near complete absence of insoluble TDP-4FL levels after the CHX recovery period. In contrast, the addition of CB5083 during the CHX recovery period resulted residual amounts of insoluble TDP-4FL protein, consistent with our hypothesis that VCP helps clear insoluble intranuclear TDP-4FL aggregates (P < 0.0001; Tukey’s post-hoc analysis P <0.01, P <0.0001; Figure 4C, 4E).

**Figure 4:**
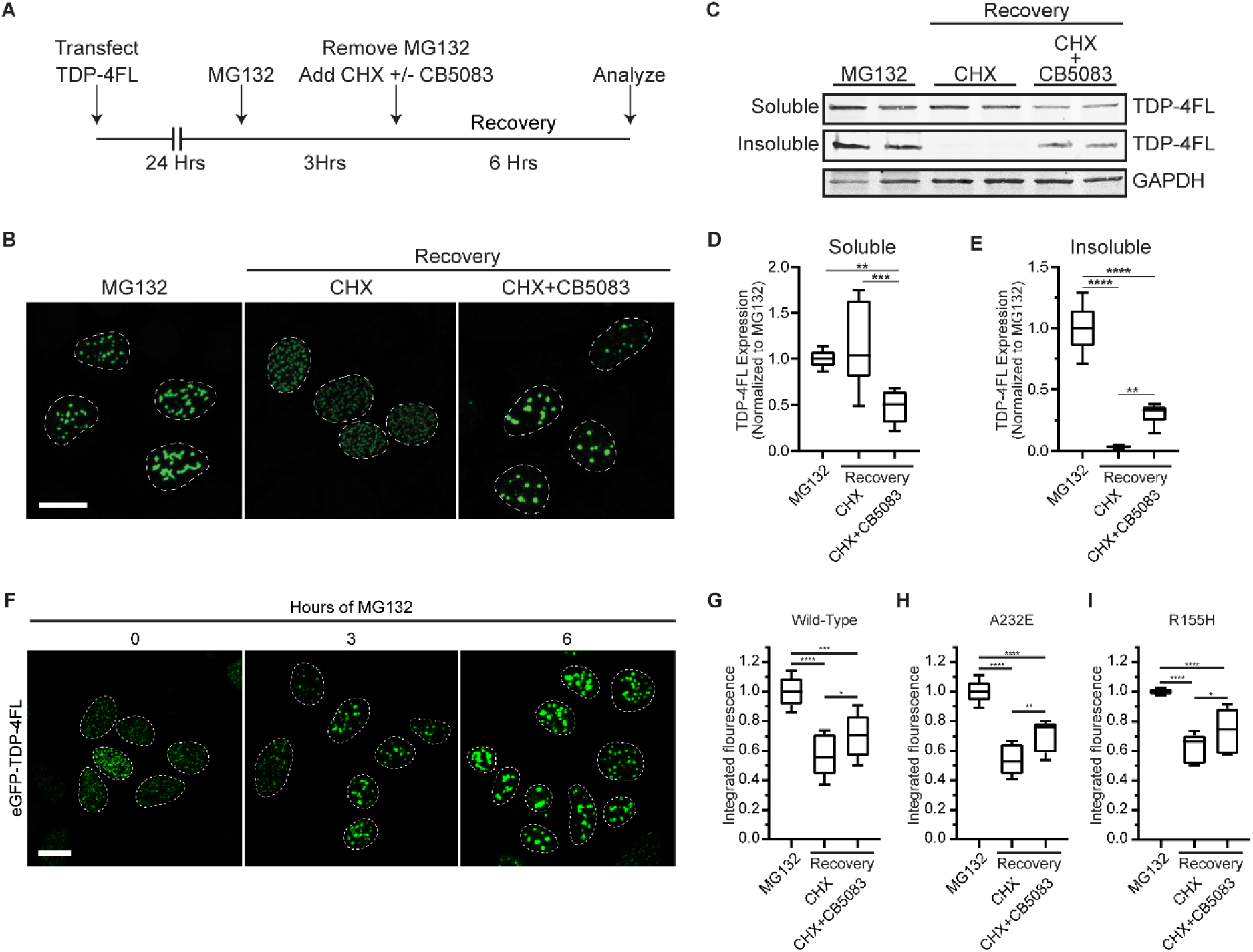
VCP inhibition impairs turnover of insoluble intranuclear TDP-4FL inclusions. **(A)** Schematic of transfection and drug treatments timing to examine the turnover of intranuclear TDP-4FL inclusions. **(B)** Representative confocal images of immunofluorescence for TDP-4FL in HeLa cells treated with 4µM MG132 for 3 hours (left), versus cells treated with MG132 for 3 hours followed by 30µg/µL CHX for 6 hours (middle) or cells treated with MG132 followed by 30µg/µL CHX + 4µM CB5083 for 6 hours (right). Scale bar, 10µm. **(C)** Immunoblots for TDP-4FL of soluble and insoluble protein fractions from HeLa cells treated with MG132 versus HeLa cells that recovered with CHX alone or CHX + CB5083. GAPDH shown as a loading control. **(D)** Quantification of soluble TDP-4FL immunoblots (n = 3 with 2 technical replicates per experiment; data shown as box-and-whisker plots, linear mixed effects model: drug treatment P < 0.0001; Tukey’s post-hoc analysis: ** P <0.01, *** P <0.001). **(E)** Quantification of insoluble TDP-4FL immunoblots (n = 3 with 2 technical replicates per experiment; data shown as box- and-whisker plots, linear mixed effects model: drug treatment P < 0.0001; Tukey’s post-hoc analysis: ** P <0.01, **** P <0.0001). **(F)** Representative confocal images of HeLa cells expressing eGFP-TDP4FL treated with 0, 3, or 6 hours of 4µM MG132. Scale bar, 10µm. **(G)** Wild-type (WT) HeLa cells or CRISPR-Cas9 knock-in HeLa cells harboring **(H)** A232E and **(I)** R155H VCP pathogenic variants cells expressing GFP-TDP-4FL were treated with 4µM MG132 for 3 hours, versus cells treated with MG132 for 3 hours followed by 30µg/µL CHX for 6 hours or cells treated with MG132 followed by 30µg/µL CHX + 4µM CB5083 for 6 hours and analyzed by flow cytometry (n = 4 experiments each with 2 technical replicates per condition, integrated fluorescence intensity shown as box-and-whisker plots, linear mixed effects model, drug treatment P < 0.0001; Tukey’s post-hoc analysis, *P < 0.05, **P < 0.01, ***P < 0.001, ****P < 0.0001).

We sought to recapitulate these findings using a quantitative live cell approach. Cells were transfected with a TDP-4FL construct containing a GFP tag, and flow cytometry was used to determine integrated fluorescence intensity (percent GFP multiplied by median GFP fluorescence) as a cumulative measurement of TDP-4FL protein levels where higher integrated fluorescence intensity was used as a measure of decreased clearance of intranuclear TDP-4FL aggregates. To confirm that GFP-TDP-4FL protein responded to proteasome inhibition similarly to what was observed above using non-GFP tagged TDP-4FL protein, cells expressing GFP-TDP-4FL were treated with MG132 for 0, 3, or 6 hours. Confocal immunofluorescence revealed that similar to myc-TDP-4FL, GFP-TDP-4FL formed intranuclear anisosome inclusions, and longer MG132 treatment led to an accumulation of larger GFP-TDP-4FL aggregates (Figure 4F). Thus, to study clearance and turnover of intranuclear GFP-TDP-4FL, integrated fluorescence intensity was measured using flow cytometry using the same experimental paradigm shown in Figure 4A. HeLa cells expressing GFP-TDP-4FL were first treated with MG132 for 3 hours to build up insoluble GFP-TDP-4FL aggregates followed by MG132 removal and recovery in CHX with or without CB5083 for 6 hours. In WT cells, recovery with CHX alone led to a decrease in integrated fluorescence intensity, consistent with the clearance of GFP-TDP-4FL aggregates upon removal of MG132. In contrast, with the presence of both CHX and CB5083 during recovery, integrated fluorescence intensity was significantly higher than CHX only treatment, indicating that CB5083 inhibited the clearance of GFP-TDP-4FL aggregates (P < 0.0001; Tukey’s post-hoc analysis P < 0.05, P < 0.001, P < 0.0001; Figure 4G). These results recapitulated our biochemical findings demonstrating that CB5083 treatment leads to decreased clearance of insoluble TDP-4FL aggregates (Figure 4C, 4E) and confirm that VCP is involved in GFP-TDP-4FL clearance. In addition, when using CRISPR edited cells expressing VCP-A232E and R155H, CB5083 also led to a greater integrated fluorescence intensity compared to CHX only treatment (P < 0.0001; Tukey’s post-hoc analysis P < 0.05, P < 0.01, P < 0.0001; Figure 4H, 4I). These findings indicate that, in the setting of pathogenic VCP variants, pharmacologic VCP inhibition further inhibits the turnover of intranuclear GFP-TDP-4FL aggregates.

### Small Molecule VCP Activators

Together, the above evidence suggests that pathogenic VCP variants are associated with a common pathology (intranuclear inclusions) and a reduction in nuclear VCP localization. Moreover, VCP activity appears to enhance clearance of intranuclear aggregates. These results suggested enhancing VCP function may counter the deleterious effects of pathogenic VCP variants. We identified four compounds, designated UP12, UP109, UP158, and UP163, that significantly increased recombinant WT VCP ATPase activity using a luciferase-based Kinase-Glo assay. Importantly, these compounds did not inhibit luciferase activity (Figure 5A, Supplemental Figure 2). These results were verified using an orthogonal ATPase assay using MESG-PNP to measure phosphate release (Figure 5B).

**Figure 5.**
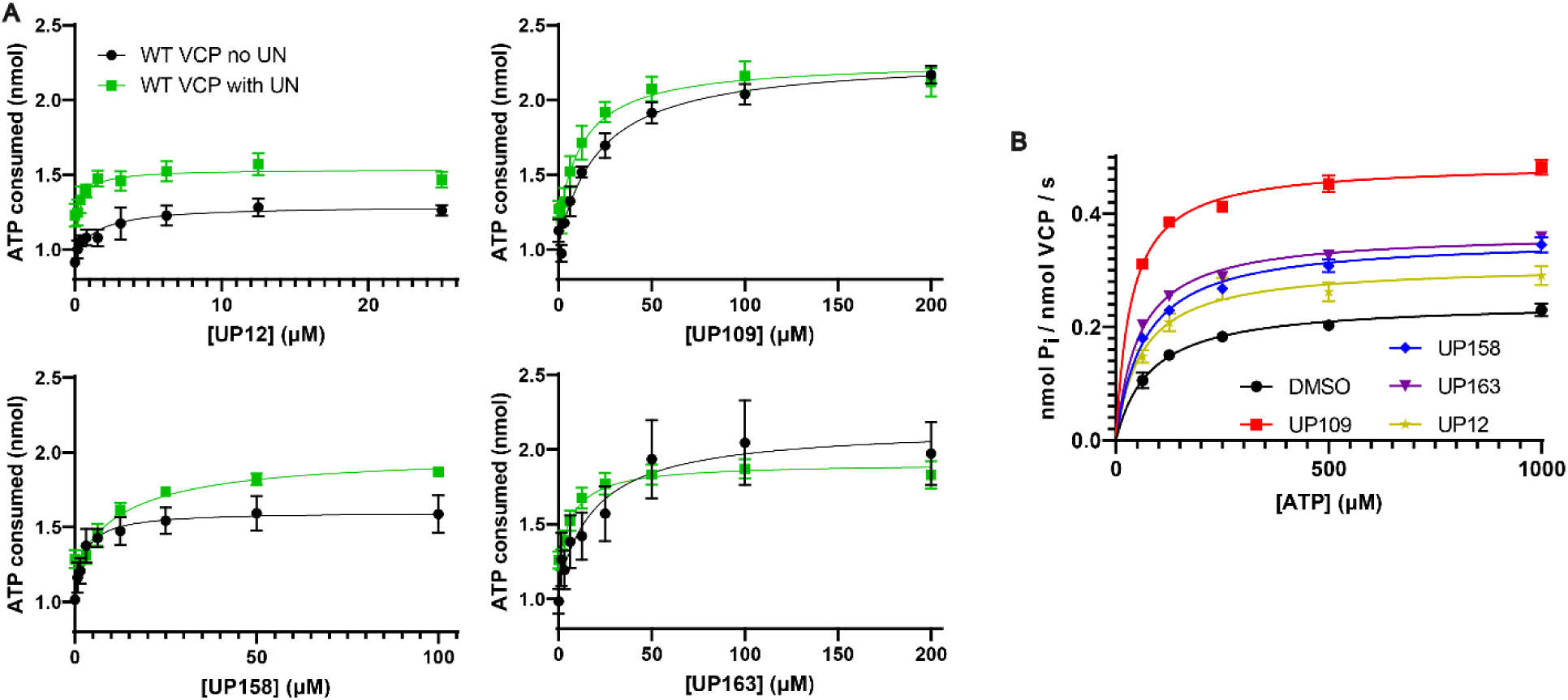
Four compounds increase VCP ATPase activity in vitro. **(A)** ATPase activity based on a Kinase-Glo assay of 50 nM recombinant VCP and 62.5 µM ATP without (black) or with 50 nM recombinant UFD1/NPLOC4 (green) versus increasing concentrations of activator. Data were fit to 3-parameter dose curve equation using least squares regression (n=3 with 3 technical replicates per experiment; points are expressed as mean ± SEM). **(B)** ATPase activity of 50 nM recombinant VCP protein was measured using an MESG-PNP assay upon treatment with 25 μM compound versus 2% DMSO control over a range of ATP concentrations. Data were plotted as v0/ET vs. [ATP] to fit a Michaelis-Menten model using least squares regression for each activator (n=3 with 3 technical replicates per experiment; points are expressed as mean ± SEM).

UP12 and UP158 exhibited a modest maximal increase of VCP activity (39% and 50% increase respectively), while UP109 and UP163 increase VCP activity 97% and 104% (Figure 5A, Table 1). UP12 and UP158 had similar EC_50_ values of 1.24 and 2.57 μM respectively, while UP109 and UP163 had relatively high EC_50_ values of 24.7 and 9.00 μM respectively. Notably, the relatively high EC_50_ for UP109 and UP163 is in part a reflection of their higher maximum activity in terms of percent ATPase activity and not necessarily an indication of lower potency.

**Table 1.** Kinetic parameters of identified compounds.

| <u>Protein</u> | WT VCP |  | WT VCP + Ufd1/Nploc4 |  | WT VCP |  |  |
| --- | --- | --- | --- | --- | --- | --- | --- |
| <u>Compound</u> | <u>EC<sub>50</sub> (μM) (95% CI)</u> | <u>Maximum % Activity (95% CI)</u> | <u>EC<sub>50</sub> (μM) (95% CI)</u> | <u>Maximum % Activity (95% CI)</u> | <u>K<sub>M</sub> (μM) (95% CI)</u> | <u>k<sub>cat</sub> (nmol P<sub>i</sub> / nmol VCP / s) (95% CI)</u> | <u>k<sub>cat</sub>/K<sub>M</sub> (s<sup>-1</sup> mM<sup>-1</sup>)</u> |
| DMSO | N/A | N/A | N/A | N/A | 80.95 (60.19, 106.6) | 0.244 (0.227, 0.261) | 3.01 |
| UP12 | 1.57 (0.51, 5.00) | 139.0 (131.9, 149.0) | 0.648 (0.240, 2.01) | 124.6 (119.8, 130.2) | 61.46 (29.70, 109.4) | 0.309 (0.269, 0.356) | 5.03 |
| UP109 | 24.72 (18.20, 34.16) | 197.4 (189.8, 206.1) | 13.36 (7.44, 25.07) | 175.3 (164.6, 188.0) | 36.17 (23.88, 50.96) | 0.489 (0.459, 0.520) | 13.52 |
| UP158 | 2.41 (1.59, 3.75) | 150.3 (146.6, 154.3) | 14.30 (7.68, 28.82) | 152.1 (143.2, 164.4) | 67.59 (40.63, 104.8) | 0.356 (0.320, 0.396) | 5.26 |
| UP163 | 9.00 (4.38, 18.94) | 204.0 (189.2, 223.3) | 8.90 (5.07, 16.60) | 151.8 (144.7, 160.7) | 54.18 (33.48, 81.35) | 0.366 (0.335, 0.401) | 6.76 |

VCP ATPase activity can potentially be enhanced by increasing the turnover rate of ATP, increasing the affinity of ATP for VCP, or a combination of the two. To determine how the activators affected k_cat_ and K_M_ of WT VCP, recombinant VCP with 25 μM activator was assessed for ATPase activity with increasing concentrations of ATP. The k_cat_, indicating the turnover rate of ATP by VCP, was significantly increased in all activator experiments from 0.244 nmol P_i_/nmol VCP/s in the DMSO condition to 0.309 with UP12, 0.489 with UP109, 0.356 with UP158, and 0.366 with UP163. UP12 had the smallest increase of about 25%, while UP158 and UP163 had a more than 50% increase and UP109 had a 100% increase in k_cat_ (Figure 5B, Table 1). The K_M_, the concentration of ATP that yields half the maximum rate of ATP turnover by VCP, was only significantly reduced with UP109 which resulted in a more than two-fold decrease in K_M_ (36.17 µM) compared to DMSO (80.95 µM) (Figure 5B, Table 1). Additionally, the ratio of k_cat_ to K_M_ can be used to determine catalytic efficiency of VCP for ATP. The catalytic efficiency of VCP with DMSO was 3.01 mM^-1^ s^-1^. All compounds increased the k_cat_/K_M_, indicating an improvement in catalytic efficiency. UP12, UP158, and UP163 were similar at 5.03, 5.26, and 6.76 respectively, while UP109 had a more than four-fold increase at 13.52 (Table 1). In the case of UP109, a synergistic effect of VCP turnover rate and affinity for ATP is indicated by the more than four-fold increase in catalytic efficiency, suggesting it may be the most effective of the four compounds described.

In vivo, protein cofactors are often required for optimal VCP activity. For example, UFD1 and NPLOC4 (“UN” collectively), allow VCP to recognize ubiquitinated substrates for unfolding upstream of the 26S proteasome. Moreover, VCP ATPase activity in vitro increases in the presence of the UN cofactors (10). To determine how this series of activators affected recombinant VCP in the presence of UN, recombinant UN was used in a 1:1 molar ratio with VCP. Compared to without UN, UP163 had no change in EC_50_, while the EC_50_ of UP109 and UP12 exhibited non-significant decreases from 24.72 µM to 13.36 µM and 1.57 µM to 0.648 µM respectively. The EC_50_ of UP158 significantly increased from 2.41 µM to 14.30 µM (Figure 5A, Table 1). Maximum percent activity stayed the same for UP158 (150.3 to 152.1) with UN compared to without UN, but significantly decreased for UP12 (139.0 to 124.6), UP109 (197.4 to 175.3), and UP163 (204.0 to 151.8) (Figure 5A, Table 1). With UN, UP12 had the lowest maximum percent activity, UP158 and UP163 had similar intermediate activities, and UP109 had the highest maximum percent activity. The varied effect of recombinant UN on EC_50_ and maximum percent activity with recombinant VCP may indicate distinct mechanisms of action represented by the four compounds.

VCP is a member of the AAA+ ATPase family of proteins that includes many proteins with similar activities in mammals and other organisms. To determine the specificity of our compounds for VCP compared to a similar human AAA+ ATPase, recombinant N-ethylmaleimide-sensitive factor (NSF) was purified. NSF ATPase activity was determined using 25 µM activator or 2% DMSO. No compound significantly increased ATPase activity of recombinant NSF compared to DMSO, indicating that UP12, UP109, UP158, and UP163 have at least some specificity for VCP over other AAA+ ATPases (Figure 6A).

**Figure 6.**
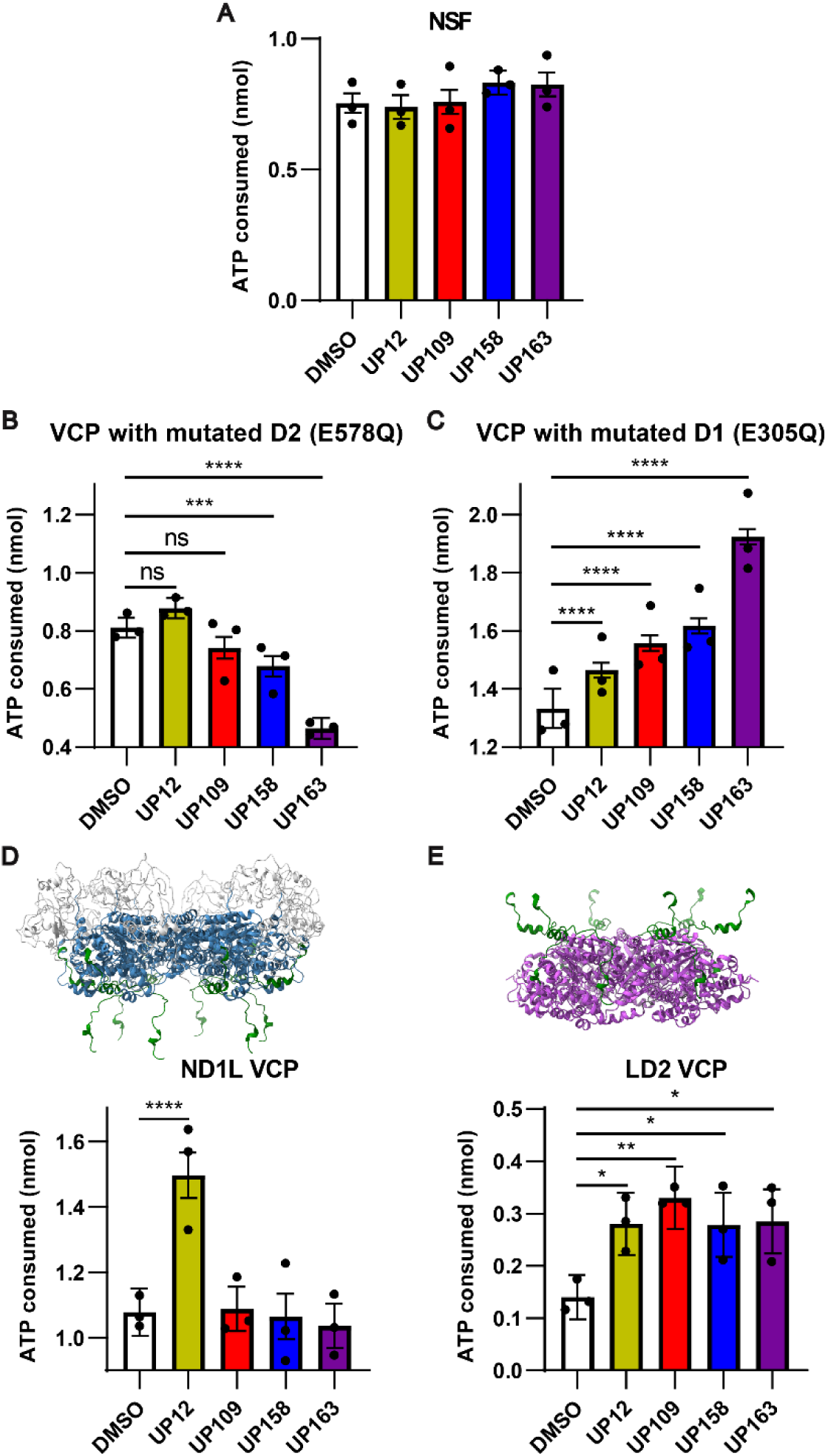
VCP activators exhibit ATPase domain specific effects. **(A)** ATP consumed as determined by a Kinase-Glo assay of 62.5μM ATP, 25μM compound or 2% DMSO, and 50nM recombinant **(A)** N-ethylmaleimide-sensitive-factor (NSF), **(B)** VCP (E578Q), **(C)** VCP (E305Q), **(D)** ND1L (aa1-481) VCP, or **(E)** LD2 (aa442-806) VCP (for all panels: n=3 with 3 technical replicates per experiment, points are expressed as beta ± SE, linear mixed effects model: *P<0.05,**P<0.01; ***P<0.001; ****P<0.0001). **(D)** and **(E)** show schematics of the VCP truncations used with N-terminal domain (White), D1 ATPase domain (blue), linker region shared between ND1L and LD2 (aa442-481) (green), and D2 ATPase domain (purple) (PDB:5FTN).

Similarly, it was important to determine the VCP ATPase domain specificity of our compounds. VCP has two distinct ATPase domains, D1 and D2, that have distinct functions. The D1 ATPase domain sits above the D2 ATPase domain in the homohexameric VCP double-ring structure. It has been suggested that ATP binding to D1 assists with initiation of substrate unfolding. D2, however, works to continuously unfold a protein substrate once it has been loaded (8,10). Walker B mutations, which allow ATP to bind, but not be hydrolyzed, were added to the D1 (E305Q) or D2 (E578Q) ATPase active sites in recombinant VCP to isolate D2 and D1 ATPase activity respectively in the full-length recombinant VCP. ATPase activity was determined with 25 µM activator or DMSO to understand ATPase domain specificity of the compounds. With an inactive D2 and active D1 (VCP E578Q), only UP12 seemed to increase ATPase activity, though not significantly (P=0.069). UP158 and UP163, interestingly, significantly decreased ATPase activity of the D1 ATPase domain (P<0.001 and P<0.0001 respectively). UP109 also decreased ATPase activity, though less than UP158 and UP163 and not significantly (P=0.074; Figure 6B). However, with an inactive D1 and active D2 (VCP E305Q) all compounds significantly increased D2 ATPase activity (P<0.0001 for all; Figure 6C). These are the first compounds to our knowledge that can increase D2 ATPase activity of VCP. UP12, uniquely, is the only compound to increase ATPase activity of both domains.

As an AAA+ ATPase and hexamer with a total of 12 ATPase domains, VCP has many allosteric interactions that coordinate activity of D1 and D2 ATPase domains. To understand how the compounds specifically affect the D1 and D2 ATPase domains in the absence of this coordination between the two ATPase domains, two truncations that isolate the D1 and D2 ATPase domain were purified. One construct (AA 1-481), termed ND1L, contained the N-terminal domain, D1, and a linker region (Figure 6D). The other construct (AA 442-806), termed LD2, contained a linker region, D2, and the C-terminus (Figure 6E). ATPase activity was again measured with 25 µM compound or DMSO. With the ND1L construct, UP12 was the only compound to increase ATPase activity (P<0.0001; Figure 6D), reminiscent of what was observed above using the full length VCP with a D2 Walker B mutation protein, VCP E578Q. Moreover, UP109, UP158, and UP163 did not inhibit the ATPase activity of the ND1L construct. The lack of inhibition with the truncated VCP compared to the full-length VCP E578Q likely indicates that UP109, UP158, and UP163 may bind outside of AA 1-481, and that the inhibition observed above with the full length D2 Walker B mutation protein (VCP E578Q) was an allosteric effect sourcing from within the D2 ATPase domain. With the LD2 construct, all compounds significantly increased ATPase activity, reminiscent of what was observed using full length VCP protein harboring the E305Q D1 domain Walker B mutation (P<0.05 for UP12, UP158, UP163; P<0.01 for UP109) (Figure 6E). UP12, as the only compound to increase ATPase activity of both ND1L and LD2 likely binds to the overlap linker region present in both truncated proteins (AA 442-481) which is known to be required for ATPase activity of the truncated VCP proteins(35–37). In contrast, UP109, UP158, and UP163 likely bind in the D2 ATPase domain.

### Clearance of insoluble intranuclear TDP-4FL aggregates is enhanced by UP109

Because UP109 modulated the greatest increase in VCP ATPase activity in our in vitro studies, we hypothesized that UP109 would enhance clearance of TDP-4FL aggregates in our cell models. To test whether UP109 would enhance clearance of TDP-4FL aggregates in our cell models, cells expressing GFP-TDP-4FL were first treated with a long MG132 treatment for 17 hours to build up insoluble intranuclear aggregates followed by removal of MG132 and addition of CHX with or without 5μM or 20μM of UP109 for 4 hours (Figure 7A). Flow cytometry analysis after 4 hours of UP109 treatment revealed that integrated fluorescence intensity was significantly decreased with 5μM of UP109, and was further reduced with 20μM of UP109 (P < 0.05, P < 0.01; Figure 7B). To exclude any potential effect of the GFP tag used in these experiments, additional experiments were conducted using a non-GFP tagged TDP-4FL. Following the same timing regiment as described above (Figure 7A), cells expressing myc-TDP-4FL were subject to MG132 treatment followed by removal of MG132 and addition of CHX with or without 5μM of UP109. Cells treated with only CHX exhibited large and dense TDP-4FL aggregates observed by confocal immunofluorescence. However, in the presence of UP109, TDP-4FL aggregate appeared to be smaller and in some cases revert to small anisosomes (Figure 7C). This was also confirmed with biochemical analysis of soluble and insoluble protein fractions collected after CHX versus CHX+UP109 treatment. Immunoblotting for TDP-4FL demonstrated that the addition of 5μM UP109 resulted in an increase in soluble TDP-4FL levels (Figure 7D, 7E) and a significant decrease in insoluble TDP-4FL levels compared to cells treated only with CHX (P < 0.05; Figure 7D, 7F). Collectively, these findings suggest that activation of VCP activity with UP109 enhances clearance of intranuclear TDP-4FL inclusions.

**Figure 7:**
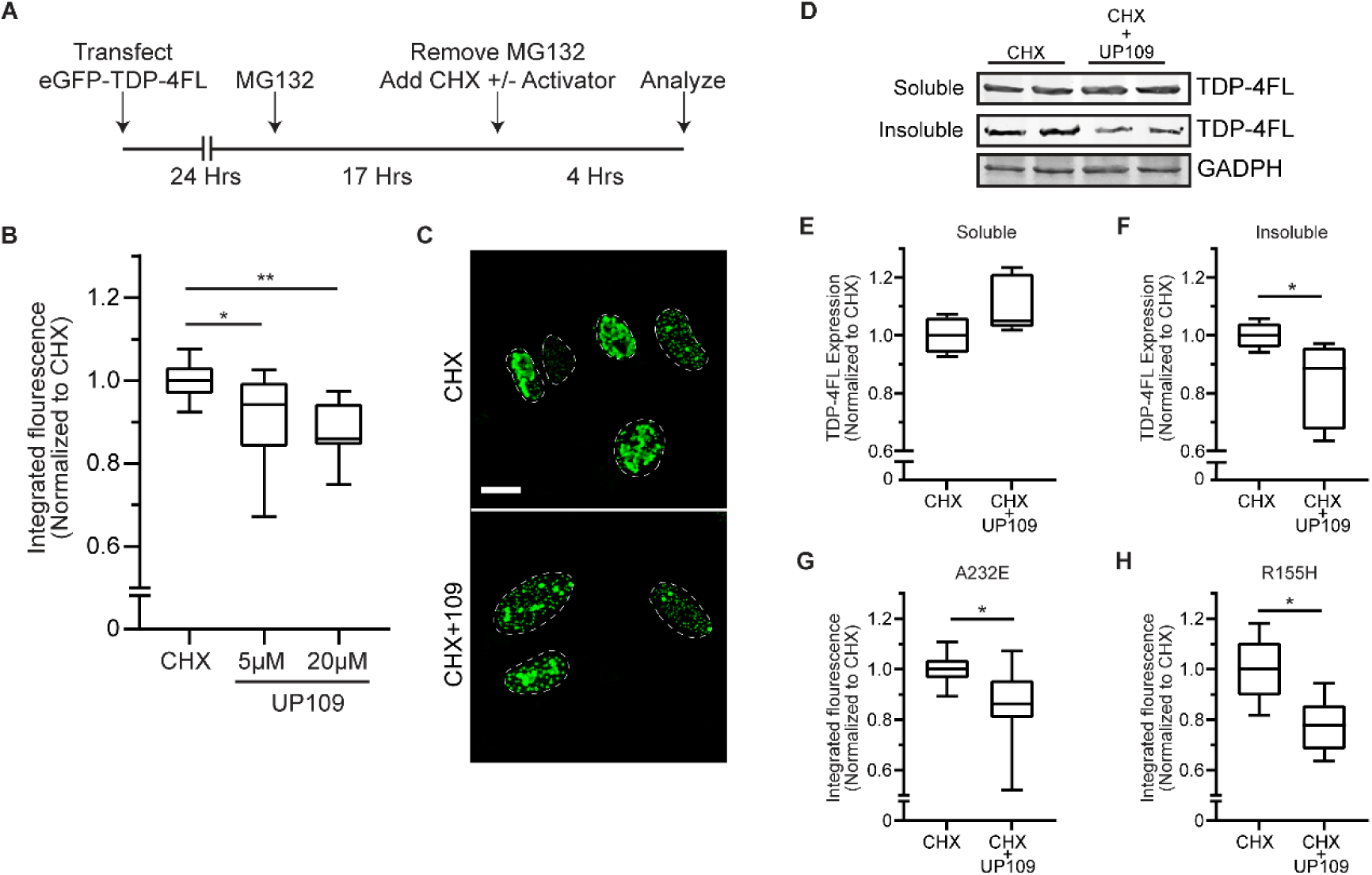
VCP activator enhances clearance of intranuclear TDP-4FL inclusions. **(A)** Schematic of transfection and drug treatments timing to examine the turnover of intranuclear TDP-4FL inclusions. **(B)** Cells expressing GFP-TDP-4FL were treated with 4µM MG132 followed by 30 µg/µL CHX with or without 5 or 20µM of activator UP109 and analyzed by flow cytometry (n = 4 experiments each with 2 technical replicates per condition, integrated fluorescence intensity shown as box-and-whisker plots, linear mixed effects model: *P < 0.05, **P < 0.01). **(C)** Representative confocal microscopy images of HeLa cells expressing TDP-4FL after treatment with 4µM MG132 followed by 4 hours of 30µg/µL CHX with or without 5µM UP109. Scale bar, 10µm. **(D)** Immunoblot for myc (TDP-4FL) of soluble and insoluble protein fractions after treatment with MG132 followed by 4 hours of 30µg/µL CHX with or without 5µM UP109. GAPDH shown as a loading control. **(E-F)** Quantification of TDP-4FL immunoblots from (E) soluble protein fractions (n = 3 experiments each with 2 technical replicates per condition; data shown as box-and-whisker plots, linear mixed effects model, P = 0.0541) and (F) insoluble protein fractions (n = 3 experiments with 2 technical replicates per condition; data shown as box-and-whisker plots, linear mixed effects model *P < 0.05). **(G-H)** VCP pathogenic variant harboring (G) A232E and (H) R155H cells were transfected with GFP-TDP-4FL and then treated with MG132 followed by 30µg/µL CHX with or without 5µM of UP109. Integrated fluorescence intensity was measured by flow cytometry (n = 4 experiments each with 2 replicate replicates per condition, data shown as box-and-whisker plots, linear mixed effects model, *P < 0.05).

Lastly, because accumulation of insoluble intranuclear TDP-4FL was exacerbated in cells harboring A232E and R155H VCP variants (Figure 3), we hypothesized that UP109 would enhance the clearance of insoluble TDP-4FL aggregates in the background of the pathogenic VCP variants. To test this, GFP-TDP-4FL was expressed in VCP-A232E and R155H cell lines and integrated fluorescence intensity was measured using flow cytometry following the same experimental paradigm shown in Figure 7A. In both cell lines, 5μM of UP109 significantly decreased integrated fluorescence intensity compared to the CHX only treatment condition (P<0.05; Figure 7G, 7H), suggesting that UP109 can enhance clearance of GFP-TDP-4FL aggregates in the setting of pathogenic A232E and R155H VCP variants.

## Discussion

In this study, we explore VCP’s role in nuclear protein homeostasis and propose a novel unifying hypothesis that MSP pathophysiology is linked to a loss of nuclear VCP function resulting in impaired nuclear proteostasis. While there is growing research in understanding how VCP variants alter cellular function, there has been limited knowledge of why VCP variants can affect seemingly distinct organ systems in MSP. Here, we highlight evidence of a common pathologic link between these tissues, showing that in addition to neurons in FTLD-TDP type D and myocytes in inclusion body myopathy, Paget’s disease of bone tissue contain osteoclasts with ubiquitinated intranuclear inclusions. The accumulation of ubiquitinated nuclear inclusions suggested that these diverse diseases may each be associated with a defect in nuclear protein homeostasis. Indeed, MSP affects cell types with a high nuclear proteostasis burden due to either multinucleation (myocytes, osteoclasts) or due to being post-mitotic where proteotoxic load cannot be diluted to daughter cells upon cell division (neurons).

Previous studies have shown that over-expression of pathogenic MSP VCP variants was associated with reduced nuclear VCP expression (29,30). Comparably, we show that knock-in of A232E or R155H into HeLa cells also resulted in a reduction in nuclear VCP expression levels. The mechanism of VCP nuclear transport has not been clearly defined. Most MSP variants are located between the N-terminal domain and D1 cleft of VCP (19). Insight to how this region may affect shuttling has been shown in previous studies where cleavage of the N-terminal domain led to a reduction in nuclear VCP expression (29,38). Additionally, VCP may contain a nuclear localization sequence within the N-terminal domain (38). How MSP variants affect nuclear localization remains to be explored. There have been some structural and biochemical studies that suggest VCP with MSP variants prefer the “up” conformation of VCP’s NTD (19) that would obscure the predicted nuclear localization sequence. However, it is difficult to determine if this potential nuclear localization sequence regulates nuclear localization as it is located at the critical junction between the N-terminus and D1 domains of VCP where mutation of this sequence is likely to have strong effects on VCP’s overall structure and function (19). In addition, post-translational modification including phosphorylation of VCP has been reported to regulate VCP nuclear localization, indicating that the mechanisms that regulate VCP subcellular localization are likely complex (39–42).

VCP’s role in the nucleus has been mostly focused on the role of the yeast VCP homologue, Cdc48, in modulating homeostasis of proteins involved in DNA replication or DNA damage (43–45). However, one study in yeast showed that Cdc48 is important for nuclear protein quality control by facilitating degradation of highly insoluble substrates downstream of nuclear ubiquitin-protein ligase, San1 (24). Many MSP in vivo models have looked at VCP-dependent defects of mitochondria function, autophagy, and endolysosomal processing; however, few have explored VCP’s role in nuclear protein homeostasis. Earlier MSP mouse models of transgenic expression of MSP variants recapitulated multiple MSP disease phenotypes and also showed some evidence of cytoplasmic TDP-43 pathology in brain (46). However, VCP variants that cause MSP leads specifically to FTLD-TDP Type D where TDP-43 inclusions are primarily localized within nuclei, which is a phenotype that has not been seen in mouse models. Here, we developed a cell model of intranuclear TDP-43 inclusions using expression of a RNA binding deficient construct of TDP-43 (TDP-4FL) to parse out VCP’s role in nuclear protein homeostasis. In cells expressing TDP-4FL, proteasome inhibition led to larger TDP-4FL aggregates and decreased TDP-4FL solubility. This phenotype was exacerbated in A232E and R155H cells, suggesting that these MSP variants were associated with a loss of nuclear VCP function that resulted in an accumulation of insoluble TDP-4FL aggregates.

Understanding whether pathogenic VCP variants cause disease due to a gain or loss of function is crucial to developing therapies to target disease. In vitro studies have shown that most MSP variants lead to an increase in VCP’s ATPase activity in vitro, suggesting a toxic gain of function. In a drosophila model of MSP, expression of VCP variants negatively altered mitochondrial morphology which was reversed upon treatment with a VCP inhibitor, CB5083 (47). Additionally, use of CB5083 in a R155H/R155H mouse model of MSP showed improved markers of muscle pathology (48). This raises the possibility that pathogenic VCP variants may affect certain cellular phenotypes through mechanisms associated with a gain of VCP function. Conversely, conditional knock out of VCP in young mice or conditional knock in of R155C variants into neurons lead to cortical brain atrophy and the accumulation of insoluble TDP-43 (22). We posit here that disease models should mirror the underlying pathology observed in human tissues, namely the presence of ubiquitinated intranuclear inclusions. Using such a model and consistent with a loss of nuclear VCP function, our studies show that pathogenic VCP variants A232E or R155H or pharmacological inhibition of VCP with CB5083 prevented proper clearance of intranuclear TDP-4FL aggregate inclusions. This result raises the possibility that VCP inhibition may be deleterious in the setting of MSP, and that increasing VCP activity could be therapeutically beneficial.

While many VCP inhibitors have been identified for various therapeutic purposes, identification of VCP activators has been limited. We identified four novel activators that have distinct mechanisms of action from two previously published compounds that moderately increase D1 ATPase activity only (49,50). The compounds identified here all increase D2 ATPase activity, with one compound, UP12, increasing both D1 and D2 ATPase activity. The exact roles of the D1 and D2 ATPase domain are not fully understood, however in vitro data would suggest that the D1 ATPase domain needs to bind, but not hydrolyze ATP for unfoldase activity, while the D2 domain’s ATPase activity is required for unfoldase activity of VCP (8,10). We show that the most active compound, UP109, can reduce insoluble nuclear aggregates in wild-type and mutant cells by boosting the clearance of nuclear TDP-43 aggregates. Our data combined with previous work (49,50) suggests that there may be multiple mechanisms of action by which small molecule activators can yield beneficial activation of VCP in cellular systems. What remains to be understood is how these compounds affect VCP’s unfoldase activity and how they affect VCP’s structure and ATPase activity while unfolding a polypeptide. Additionally, further work should be done to characterize if the compounds’ beneficial effect is as simple as increasing VCP activity. If this is the case, since UP109 appears to improve MSP variant pathology in our cellular models, this would suggest that pathogenic MSP variants cause disease via a loss of normal nuclear function.

In summary, our study proposes a new mechanism of disease pathogenesis for MSP that potentially links pathological findings seen in brain, muscle, and brain. We highlight the importance of VCP in maintaining nuclear protein homeostasis and its role in the clearance of insoluble, nuclear TDP aggregates. Additionally, we find that disease-linked VCP variants potentiates the accumulation of TDP inclusions which can be alleviated by a novel VCP activator. Because VCP is involved in multiple cellular processes, we cannot ignore the likelihood that pathogenic VCP variants result in defects across multiple VCP-dependent protein homeostasis pathways over time. Thus, further work in understanding the degree in which VCP variants affect different disease-relevant pathways can pave way for continued development of disease modifying therapies.

## Methods

### Generation of VCP Mutant CRISPR cell lines

To generate the A232E and R155H VCP mutant cell lines, CRISPR gRNAs were designed using Benchling software (http://benchling.com/) and cloned into pSpCas9(BB)-2A-Puro (PX459) V2.0 vector (plasmid #62988, gift from Feng Zhang, Addgene, Cambridge, MA, USA) (Supplemental Table 1). Homology directed repair templates were designed to include PAM mutation sites. Templates were synthesized by Integrated DNA Technologies, Inc. (Coralville, IA, USA). PX489 and repair template were co-transfected into HeLa cells using Lipofectamine 3000 (Invitrogen, Waltham, MA, USA). On day two post-transfection, cells were selected in media containing 1.25μg/μL of puromycin for two days. After two additional days of recovery, cells were plated onto 96-well plates for clonal isolation. DNA was extracted from 96-well plates of confluent clones using QuickExtract DNA Extraction Solution (Lucigen Corporation, Middleton, WI, USA) according to manufacturer’s protocol. DNA from clones were PCR amplified and screened for homology directed repair by restriction enzyme digest. CRISRP-edited cell lines were Sanger sequenced to verify knock-in of the mutations. Additionally, to ensure that the guides were specific to the sites of interest, the top five potential off-target sites as well as targets that were in a gene encoding region were evaluated for cutting or homology directed repair. Primers were designed for each site, amplified with PCR, and analyzed by Sanger sequence analysis (Supplemental Table 1). Our analysis confirmed that there were no off-target mutations in the A232E and R155H cell lines.

### Cell culture and plasmid transfections

Wild-type and CRISPR-Cas9 knock-in HeLa cell lines harboring A232E or R155H VCP pathogenic variants were maintained at 37°C and grown in DMEM high glucose (Invitrogen) supplemented with 10% fetal bovine serum (Atlanta Biologicals, Flowery Branch, GA, USA) and 2mM L-Glutamine (Invitrogen). For transfection of CRISPR components, 1.5μg Cas9/gRNA containing plasmid and 1.5μg repair template were co-transfected into 6-well dishes using Lipofectamine 3000 (Invitrogen) with a 3:1 transfection reagent to DNA ratio for 48 hours. For transient transfections, 1.5μg of plasmid was transfected into 6-well dishes using Fugene HD (Promega, Madison, WI, USA) with 3:1 transfection reagent to DNA for 24 hours. Myc-TDP-43 and myc-TDP-4FL plasmid were gifted from the Center for Neurodegenerative Disease Research (CNDR, University of Pennsylvania, Philadelphia, PA, USA). GFP-TDP-4FL was generated by introducing F147/149/229/231L mutations using site directed mutagenesis with QuickChange XL kit according to the manufacturer’s protocol (Agilent Technologies, Santa Clara, CA) into GFP-TDP-43 plasmid (gifted from CNDR).

### Drug treatments and reagents in cell experiments

For proteasome inhibition, cells expressing myc-TDP-43 or myc-TDP-4FL were treated with 4μM of MG132 (M8699, Sigma, St. Louis, MO, USA) for 0, 3, or 6 hours. For recovery experiments, cells expressing myc-TDP-4FL or GFP-TDP-4FL were treated with 4μM of MG132 for 3 hours followed by removal of MG132 and the addition of 30μg/μL of cycloheximide, CHX (C1988, Sigma) with DMSO or 2.5μM of CB5083 (19311, Cayman Chemical Company,Ann Arbor, MI, USA) for 6 hours. To test efficacy of UP109 activator, cells expressing myc-TDP-4FL or GFP-TDP-4FL were treated with 4μM of MG132 for 17 hours followed by removal of MG132 and the addition of 30μg/μL of cycloheximide, CHX with DMSO or 5μM or 20μM of UP109 (Enamine Ltd, Kyiv, Ukraine) for 4 hours.

### Sequential protein extraction

To collect soluble and insoluble protein fractions, cells were subject to sequential extraction using buffers of increasing strengths (RIPA and then urea buffers). Cells were cultured in a six-well dish and pelleted by centrifugation. Cell pellets were washed with cold 1x DPBS and re-suspended in cold RIPA buffer containing 150mM NaCl, 1% Triton X-100, 0.5% sodium deoxycholate, 0.1% SDS and 50mM Tris (pH 8.0) supplemented with protease inhibitors. Samples were sonicated and centrifuged at 100,000 x g for 30 minutes at 4°C. Supernatant was collected as RIPA soluble fraction. Pellet was then re-extracted in RIPA buffer to remove residual soluble proteins. The insoluble pellet was extracted in 1/3 of starting volume of urea buffer containing 7M Urea, 2M Thiourea, 4% CHAPS, 30mM Tris (pH 8.5), sonicated, and centrifuged at 100,000 x g for 30 minutes at 22°C. The supernatant was collected as the RIPA insoluble (urea soluble) fraction.

### Immunoprecipitation

Myc immunoprecipitation were carried out using Anti-c-Myc Magnetic Beads (Pierce, Thermo Fisher Scientific, Waltham, MA, USA). Beads prepared following manufacturers instruction and incubated with urea soluble fractions diluted in 1x TBST Buffer (25mM Tris, 0.15M NaCl, 0.05% Tween-20 Detergent) containing 100mM N-Ethylmaleimide (NEM) and 5mM of EDTA to preserve ubiquitination of protein lysates, overnight at 4°C. To elute samples from beads, Laemmli sample buffer containing 0.5mM dithiothreitol was added directly to beads and heated for 10 minutes at 99°C.

### Western blot and antibodies

Protein concentration was determined using BCA reagent (Thermo Fisher Scientific). 25-40μg of protein was run on 7.5-15% sodium dodecyl sulfate-polyacrylamide gel electrophoresis (SDS-PAGE) gel and transferred using the Trans-Blot Turbo system (Biorad, Hercules, CA) to nitrocellulose membranes. Membranes were blocked using 5% non-fat milk and incubated in primary antibodies in 1x TBST (1x TBS + 0.1% Tween-20) overnight at 4°C. For immunoprecipitation experiments, nitrocellulose membranes were transferred using Mini Trans-Blot Cell (Biorad) overnight. IR dye secondary antibodies (Licor Biosciences, Lincoln, NE, USA) were used to detect protein using a Licor Odyssey. Primary antibodies used include mouse anti-Myc (9E10), rabbit anti-Myc (ab9106, Abcam, Boston, MA), rabbit anti-TDP-43 (C2089, gift from CNDR), mouse anti-VCP (NB120-11433, Novus Biologicals, Centennial, CO, USA), rabbit anti-ubiquitin (43124, Cell Signaling Technology, Danvers, MA), and rabbit anti-GAPDH (2118, Cell Signaling Technology).

### Immunofluorescence

HeLa Cells were plated onto glass coverslips and transfected with TDP-4FL using Fugene HD (Promega) for 24 Hours. Cells were fixed with 4% paraformaldehyde (Electron Microscopy Sciences, Hatfield, PA, USA), permeabilized with 0.1% Triton-X (Thermo Fisher Scientific), and blocked with 2% FBS 1x DPBS. Primary antibodies were diluted in 2% FBS in 1x DPBS. Cells were stained with 300nM of DAPI, and coverslips were mounted onto glass slides using ProLong Glass Antifade Mountant (Thermo Fisher Scientific). Confocal images were obtained using a Leica TCS SPE laser scanning confocal microscope. Primary antibodies used include mouse anti-Myc (9E10), rabbit anti-Myc (ab9106, Abcam), mouse anti-VCP (NB120-11433, Novus Biologicals), and mouse anti-ubiquitin (ST1200, Millipore, Burlington, MA, USA) with Alexa Flour 488 or 568 (Invitrogen) as secondary antibodies.

### Image analysis of nuclear to cytoplasmic fluorescence intensity ratio

Confocal images of immunofluorescence staining for VCP in wild-type or CRISPR-Cas9 knock-in HeLa cells harboring A232E and R155H VCP pathogenic were analyzed using Fiji (ImageJ). Nuclei and cytoplasmic regions were segmented and fluorescence intensity was determined for each subcellular region. Cytoplasmic VCP fluorescence intensity values were divided by nuclear VCP fluorescence intensity value for each individual cell to determine the fluorescence intensity ratio of nuclear to cytoplasmic VCP expression.

### Immunohistochemistry

Immunohistochemistry was performed on 6 µm formalin-fixed, paraffin-embedded tissue sections from FTLD-TDP type D neocortex, inclusion body myopathy muscle, Paget’s disease bone, normal control bone, giant cell tumor of the bone, or fracture callus tissue. After deparaffinization and rehydration of the tissue, sections were treated with methanol/H202 for 30 min and then washed for another 10 min. Microwave antigen retrieval in citric acid-based antigen unmasking solution (Vector Laboratories, Burlingame, CA, USA) was then performed. Sections were washed in 0.1 M Tris buffer and blocked in 2% fetal bovine serum (FBS) in 0.1 M Tris buffer. Sections were then incubated with primary antibodies overnight at 4°C in a humidified chamber. Sections were again washed in 0.1 M Tris buffer, blocked in 2% FBS in 0.1 M Tris buffer, and then incubated with biotinylated species-specific secondary antibodies for 1–2 h at room temperature. Afterwards, sections were once more washed and blocked and incubated with avidin-biotin solution (Vectastain ABC kit, Vector Laboratories) for 1.5 h at room temperature. Sections were once more washed and developed using DAB (3,3′-diaminobenzidine) peroxidase substrate kit (Vector Laboratories). Finally, sections were dehydrated in an ascending ethanol series, and cleared using xylene. Mounting media (Cytoseal TM 60, Thermo Fisher Scientific) and glass coverslips (Thermo Fisher Scientific) were used to coverslip the slides. Primary antibodies used include rat anti-phosphoTDP-43 (1D3, TDP-43 p409/410, gifted from Drs. Maneula Neumann and Elisabeth Kremmer) and mouse anti-ubiquitin (NB300-130, Novus Biological).

### Flow cytometry and data analysis

HeLa cells were collected using trypsin and pelleted at 1000xg for 3 minutes. Cell pellets were resuspended in colorless DMEM (Invitrogen) supplemented with 2% FBS and filtered through a BD tube with cell strainer (BD Biosciences, Franklin Lakes, NJ, USA) for flow cytometry analysis on LSR II (BD Biosciences). Data analysis were performed on FlowJo v10 software. Integrated fluorescence intensity was determined by multiplying percent positive GFP and GFP median fluorescence values.

### Recombinant Protein Purification

N-terminal histidine tagged wild-type VCP was obtained from Addgene (plasmid #12373, gift from Axel Brunger). N-terminal histidine tagged NSF, VCP ND1L, and VCP LD2 were purchased from Genscript in pET-28a(+) vector. QuikChange II site-directed mutagenesis kit (Agilent Technologies) was used to generate VCP point mutations. Plasmid sequences were confirmed with Sanger sequencing.

For all VCP forms and NSF, Plasmids were transformed into *Eschericia coli* BL21-CodonPlus (DE3)-RIL cells (Agilent Technologies) following manufacturer’s instructions. Cells were grown in Terrific Broth supplemented with kanamycin at 37° C and induced at OD_600_ ∼0.800 with 1mM IPTG for 18 hours at 18° C. Cells were collected via centrifugation at 4000xg for 15 minutes at 4° C, then resuspended in lysis buffer (20mM NaH_2_PO_4_ pH 7.4, 500mM NaCl, 10mM MgCl_2_, 1mM ATP, 2mM β-mercaptoethanol, 20mM imidazole, 1mg/mL lysozyme, EDTA-free Complete Protease inhibitor (Roche)) at 4° C followed by sonication. Lysates were cleared via centrifugation at 27,500xg for 30 min at 4° C. The supernatant was filtered, then applied to a 5mL HisTrap™ excel nickel column (Cytiva, Marlborough, MA, USA). The column was washed with 10 column volumes of wash buffer (20mM NaH_2_PO_4_ pH 7.4, 500mM NaCl, 10mM MgCl_2_, 1mM ATP, 2mM β-mercaptoethanol, 20mM imidazole, EDTA-free Complete Protease inhibitor (Roche, Mannheim, Germany). Proteins were eluted using a 5mL linear gradient of elution buffer 1 (20mM NaH_2_PO_4_ pH 7.4, 500mM NaCl, 10mM MgCl_2_, 20mM imidazole) to elution buffer 2 (20mM NaH_2_PO_4_ pH 7.4, 500mM NaCl, 10mM MgCl_2_, 500mM imidazole) followed by 50mL of elution buffer 2. Elute was concentrated using an Amicon Ultra-15 centrifugal unit (Millipore) then loaded onto a Superdex 200 Increase 10/300 GL column (Cytiva). 0.5mL fractions were collected in 50mM HEPES pH 7.4, 150mM KCl, 2mM MgCl_2_, 5% glycerol. Protein concentration was determined using the Peirce™ BCA protein assay (Thermo Fisher Scientific). Fractions were concentrated to 5mg/mL using an Amicon Ultra – 0.5mL centrifugal unit (Millipore), flash frozen, and stored in liquid nitrogen. Expression plasmid was isolated from each protein preparation and confirmed by Sanger sequencing. Each preparations purity was assessed by gel electrophoresis.

UFD1 and NPLOC4 were purified by MD Anderson Cancer Center Core for Biomolecular Structure and Function similarly via the following protocol. NPLOC4 untagged plasmid and His-UFD1 were expressed separately in *E. coli* (DE3) cells grown in terrific broth at 37° C until ∼OD_600_ of 1.0, then were induced with 0.4mM IPTG overnight at 16° C. Cells were harvested by centrifugation, then re-suspended in 50mM Tris pH 7.4, 500mM KCl, 5mM MgCl_2_, 20mM imidazole, 5% glycerol, and 2mM β-mercaptoethanol with complete protease inhibitors. The resuspension pellets were mixed 1:1 Ufd1:Nploc4 and lysed by sonication. The lysate was centrifuged for 1 hour to remove insoluble material. The Ufd1-Nploc4 complex was purified from lysate using Ni-NTA resin in 50mM Tris pH 8.0, 500mM KCl, 5mM MgCl_2_, 20mM imidazole, 5% glycerol, and 2mM β-mercaptoethanol. Resin was washed using the same buffer, then eluted using 300mM imidazole. The protein elute was applied to an S75 16/16 column (Cytiva) with buffer 20mM HEPES pH 7.4, 250mM KCl, 1mM MgCl_2_, 5% glycerol, and 0.5mM TCEP. Concentration of UFD1-NPLOC4 was estimated as 1mg/mL, flash frozen, and stored at −80° C. Purity was assessed via gel electrophoresis and dynamic light scattering.

### ATPase Activity Assays

ATPase activity was determined with orthogonal methods. Experiments with varied activator concentration and screening were measured using the Kinase-Glo Luminescent Kinase Assay (Promega). Reactions were incubated at 25°C for 1 hour then reduced to 4°C in a thermocycler consisting of 50nM of recombinant VCP with or without 50nM UFD1/NPLOC4 in 50uL ATPase reaction buffer (25mM HEPES, pH 7.5, 100mM NaCl, 10mM MgCl_2_) at 2% DMSO with 62.5µM ATP and a range of activator concentrations. Equal volumes of reaction and Kinase-Glo reagent were incubated for 20 min at room temperature in a 96 well white flat bottom plate (Corning, Corning, NY, USA). Luminescence was measured in a microplate reader (Spark 20M, Tecan, Männedorf, Switzerland).

All other experiments were performed using the EnzCheck phosphatase assay kit (Molecular Probes, Eugene, OR, USA). ATPase activity was measured in a 96 well flat bottom plate (Thermo Fisher Scientific). 50nM VCP was incubated in 100µL ATPase buffer with 2% DMSO with 25µM compound and various ATP concentrations. The absorbance at 360 ± 5 nm was measured for more than 15 min at 25°C. Inorganic phosphate release was quantified based on a phosphate standard curve performed on the same plate. Initial enzyme velocity was determined by fitting the linear portion of the data to a linear regression model.

Experiments with varied compound concentration were fit to a 3-parameter dose curve with asymmetric profile likelihood 95% confidence intervals in GraphPad Prism 9 (GraphPad, San Diego, CA, USA). Experiments with varied ATP concentration were fit to a Michaelis-Menten model with asymmetric profile likelihood 95% confidence intervals. Significance was determined for experiments with fixed compound or fixed ATP concentration using a linear mixed-effects model.

### Statistical Analysis

One- or two-way ANOVA statistical analysis, Dunnett’s multiple comparison post-hoc analysis were performed using GraphPad Prism 9. 3-parameter dose curve with asymmetric profile likelihood 95% confidence intervals and Michaelis-Menten model with asymmetric profile likelihood 95% confidence intervals were fit in GraphPad Prism 9. Linear mixed effects regression models were performed using nlme package and Tukey’s multiple comparison post-hoc analysis were performed using emmeans package in RStudio. Box and whiskers plot: whiskers represent minimum to maximum; line represent median, bounds represent 25th to 75th percentiles. P value <0.05 was considered significant. P values were designated as * ≤0.05, ** ≤0.01, *** ≤0.001, **** ≤0.0001.

## Acknowledgments

We thank Bernice Benoit, Katherine Xia and the Center for Neurodegenerative Disease Research for their assistance. We thank Dr. Virginia M.-Y. Lee and CNDR for sharing antibodies and plasmids, and Dr. Manuela Neumann and Dr. Elisabeth Kremmer for providing the p409-410 antibody (TAR5P-1D3).

## Funding

This study was supported by grants from the NIH RF1AG065341, P01AG066597, P30AG072979, T32AG000255, T32GM007170, T32GM132030, and F30AG077756.

## Author contributions

JMP, BCC and EBL conceived of and designed the experiments, analyzed results, and wrote the manuscript. JMP, BCC, ATN, DDB, and NFD performed the experiments. All authors edited the manuscript.

## Competing interests

A patent application related to this work is pending. The authors declare no additional competing interests.

## Data and materials availability

Requests for data or materials should be addressed to.

## Notes

Conflict-of-interest statement: A patent application related to this work is pending. The authors have declared that no additional conflict of interest exists.

### Competing Interest Statement

A patent application related to this work is pending. The authors have declared that no additional conflict of interest exists.

