## Supplemental Information for "Novel VCP activator reverses multisystem proteinopathy nuclear proteostasis defects and enhances TDP-43 aggregate clearance"

### Supplementary Figure 1

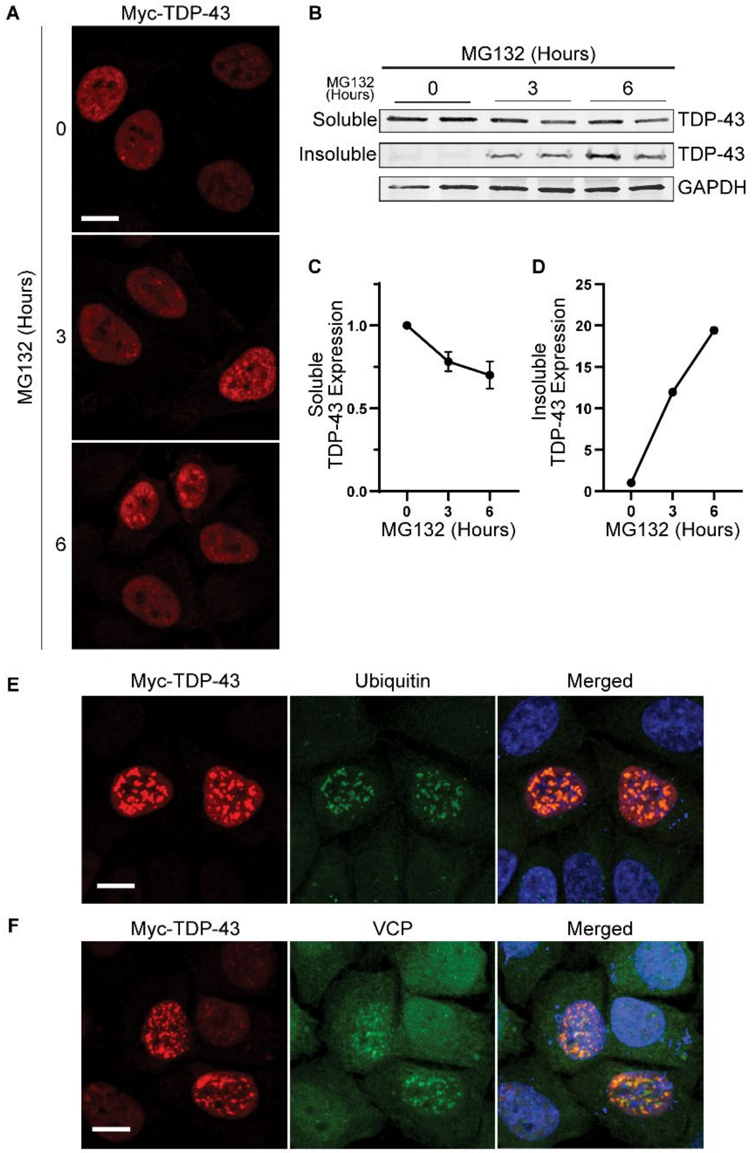


**Supplemental 1: Proteasome inhibition induces insoluble intranuclear TDP-43 inclusions that colocalize with ubiquitin and VCP.**

**(A)** Confocal images of myc (TDP-43) immunofluorescence in HeLa cells expressing myc-TDP-43 treated with 4μM MG132 for 0, 3, or 6 hours. **(B)** Immunoblot for myc (TDP-43) of soluble and insoluble protein fractions from HeLa cells expressing myc-TDP-43 with 4μM MG132 for 0, 3, or 6 hours. GAPDH shown as a loading control. Quantification of **(C)** soluble or **(D)** insoluble TDP-4FL immunoblots. (Soluble, n = 2 experiments each with 2 replicate replicates per condition, results are expressed as mean ± SEM over time. One-way ANOVA, P = 0.170 for soluble protein, *P < 0.05 for insoluble protein). **(E-F)** Immunofluorescence confocal images of HeLa cells expressing TDP-43 treated with 4uM MG132 for 3 hours stained for myc (TDP-43, red) and either ubiquitin (E, green) or VCP (F, green). Scale bar, 10μm.

#### Supplementary Figure 2

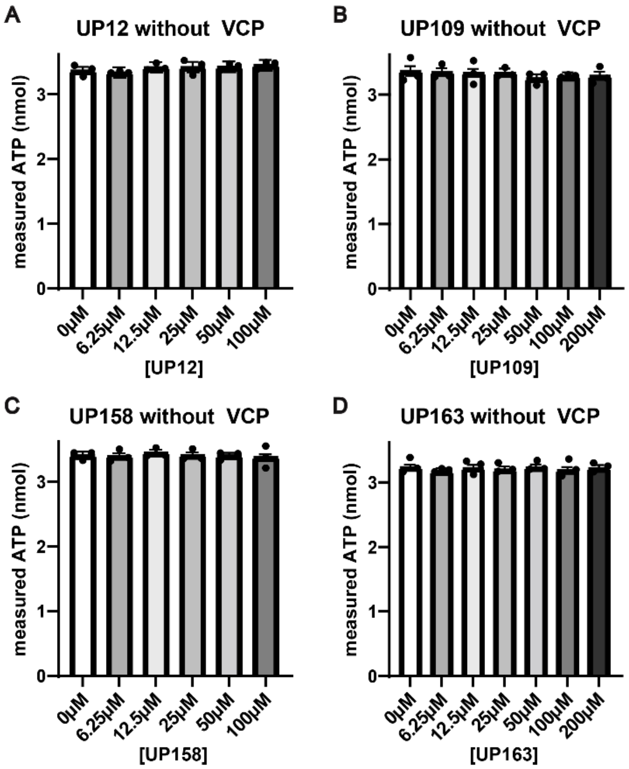


**Figure S2. VCP activator compounds do not inhibit luciferase.**
**(A-D)** Measured ATP as determined by a Kinase-Glo assay of 62.5µM ATP in the presence of increasing concentrations of compound without recombinant VCP present. (for all panels: n=3 with 3 technical replicates per experiment, points are expressed as beta ± SE, linear mixed effects model, P>0.05 for each compound).

### Supplemental Table 1.

**A232E gRNA:**

5' CACCGCCAATTGCCTTAAAGAGGGC

Primers used to amplify region off-target sequence

| Off-target Sequence | Score | Gene | Chromosome | Base Pair Mismatches | Forward Primer | Reverse Primer |
|---|---|---|---|---|---|---|
| GCACCTGCCTTAAAGAGGGC | 2.503293808 | | chrX | 3 | GCCCTCCTGCAGATTCATTG | CAGAGTTGCTTGCTTGGTGT |
| CCGTTTCCCTTAAAGAGGGC | 1.685813175 | | chr3 | 3 | CCCCACTTTTCTGCCTTTGG | CCCCTCCAAACACTGTAGCA |
| CAGATTGCCTGAAAGAGGGC | 1.627131455 | | chr7 | 3 | AAGGAAGGTGAAGGAGGAGC | TCAGTTCGCCATCTTGACCT |
| CCAAGTGGCTAAAAGAGGGC | 1.483122363 | ENSG00000166313 | chr11 | 3 | CTCAGGGTATGGGCTCTCAG | AGCTTACAGTGATGGGGCTT |
| CCGATGGCCTGAAAGAGGGC | 0.948779615 | ENSG00000261150 | chr8 | 3 | TGAGTGTCTTCATCCAGGCA | CTAAGGATGGGACCAGGGTG |
| CCAAATAGCTCAAAGAGGGC | 0.546728365 | ENSG00000171786 | chr1 | 4 | GCTCTCTCACTGAATGCTGC | ATGCAGGTTCTAAGGCCCTT |

**R155H gRNA**

5' CACCGACATTTTTCTTGTCCGTGG

Primers used to amplify region off-target sequence

| Off-target Sequence | Score | Gene | Chromosome | Base Pair Mismatches | Forward Primer | Reverse Primer |
|---|---|---|---|---|---|---|
| ACAATTTCCTTGTCCGTGGA | 1.682165605 | | chr12 | 3 | TGGCCCCATCTCTGTGAAT | TGACAGAGGCCAAGTTTTGC |
| TCATTTTCCTTGTCAGTGGT | 0.981117534 | | chr18 | 3 | AGAAGGGCCATTGGATGTCA | ACACCATTGCTCAGTAGGCT |
| CAAATTTTGTTGTCCGTGGT | 0.934200644 | | chr10 | 4 | TAGCCTCAAGTCCCAATGCA | GAGCCCAAAGTGCAGTTGAA |
| ACAGATTTCTTGTCAGTGGT | 0.861590525 | | chr9 | 3 | CCTCTCCCAGCTGTCCTTAG | ATGAAACCATGGCCCTCAGA |
| AAATATTCCTTGTCCGTGAT | 0.634004237 | | chr7 | 4 | GAAGCTGTGCAGACGGAATT | GGTCATGTGTTTGTGGCTGT |
| ACACTTTCCTTTTCCGTGGA | 0.403810359 | ENSG00000101745 | chr18 | 4 | GCAGGGATGCCAAAAGGAAA | TGGGCAGTGTTACAGGTGTA |
| TGATTTTTCTTGTCAGTGGA | 0.274572931 | ENSG00000124198 | chr20 | 4 | GGTGTCTTCTGCCCTCTCTT | AAATGGTGTCCTGGCCTGAT |
| ATTTTTTTCTTGTCAGTGGA | 0.25657315 | ENSG00000204450 | chr11 | 4 | TCTCATGGCGGCATCAGTAT | GAAACACCCCAACTTGACCC |
| ATTTTTTTCTTGTCAGTGGA | 0.25657315 | ENSG00000189253 | chr11 | 4 | AAGGTGGAGAGTTGGTGGAG | TCTCATGGCGGCATCAGTAT |
| AGATTTATTTTGTCAGTGGT | 0.196270732 | ENSG00000145495 | chr5 | 4 | CCAGACTCCTCCTTTACCCC | TCAGGAGTTCGAGACCAACC |

**Supplemental Table 1: gRNA used for A232E and R155H CRISPR editing and list of predicted off-target sites tested in CRISPR edited cells.**
